# CopY as a Transcriptional Duet Orchestrating Copper Homeostasis via Asymmetry-Dependent DNA Affinity

**DOI:** 10.64898/2026.09.30.755846

**Authors:** Melanie Hirsch, Aparna L. Benny, Lukas Hofmann, Yulia Shenberger, Lada Gevorkyan-Airapetov, Sunil Saxena, Sharon Ruthstein

**Affiliations:** Department of Chemistry and the institute of Nanotechnology and Advanced Materials, Bar-Ilan University, Ramat-Gan, Israel; Department of Chemistry, University of Pittsburgh, PA

**Keywords:** Metal transcription factor, protein-DNA interaction, EPR, DEER, RIDME, bacterial resistance

## Abstract

Copper is essential yet potentially toxic, requiring tight regulation of intracellular concentration. The copper-responsive transcriptional repressor CopY is unusual among bacterial metalloregulators because it occupies two operator sites within the *cop* promoter. Although two CopY dimers bind these operators independently, the mechanistic significance of this dual-operator architecture remains unclear. Here, we combine biochemical assays, circular dichroism (CD), and electron paramagnetic resonance (EPR) distance measurements to define the molecular basis of CopY-mediated repression. CopY binds the proximal operator with higher affinity than the distal operator. CD measurements further show that DNA binding increases CopY thermal stability, with greater stabilization by the proximal operator. Conversely, DNA-based EPR measurements reveal comparable CopY-induced compaction of both operators. Thus, protein stabilization and DNA deformation represent distinct components of repression. Moreover, site-directed spin labeling and EPR demonstrate that the metal binding domain is conformationally heterogeneous and highly flexible. DNA binding has only a marginal effect on this flexibility. Thus, the metal site is not a rigid pocket but a dynamic one, even in the DNA-bound repressor, a property that may facilitate Cu(I) capture and hence derepression once copper levels rise. We propose that the proximal operator serves as a high-affinity anchoring site, whereas the distal operator primarily promotes promoter remodeling, and that the plasticity of the metal binding domain is retained in the DNA-bound repressor. Together, these findings support a “transcriptional duet” model in which two CopY dimers bound to non-equivalent operators make distinct and complementary contributions to repression.

**Graphical Abstract:** 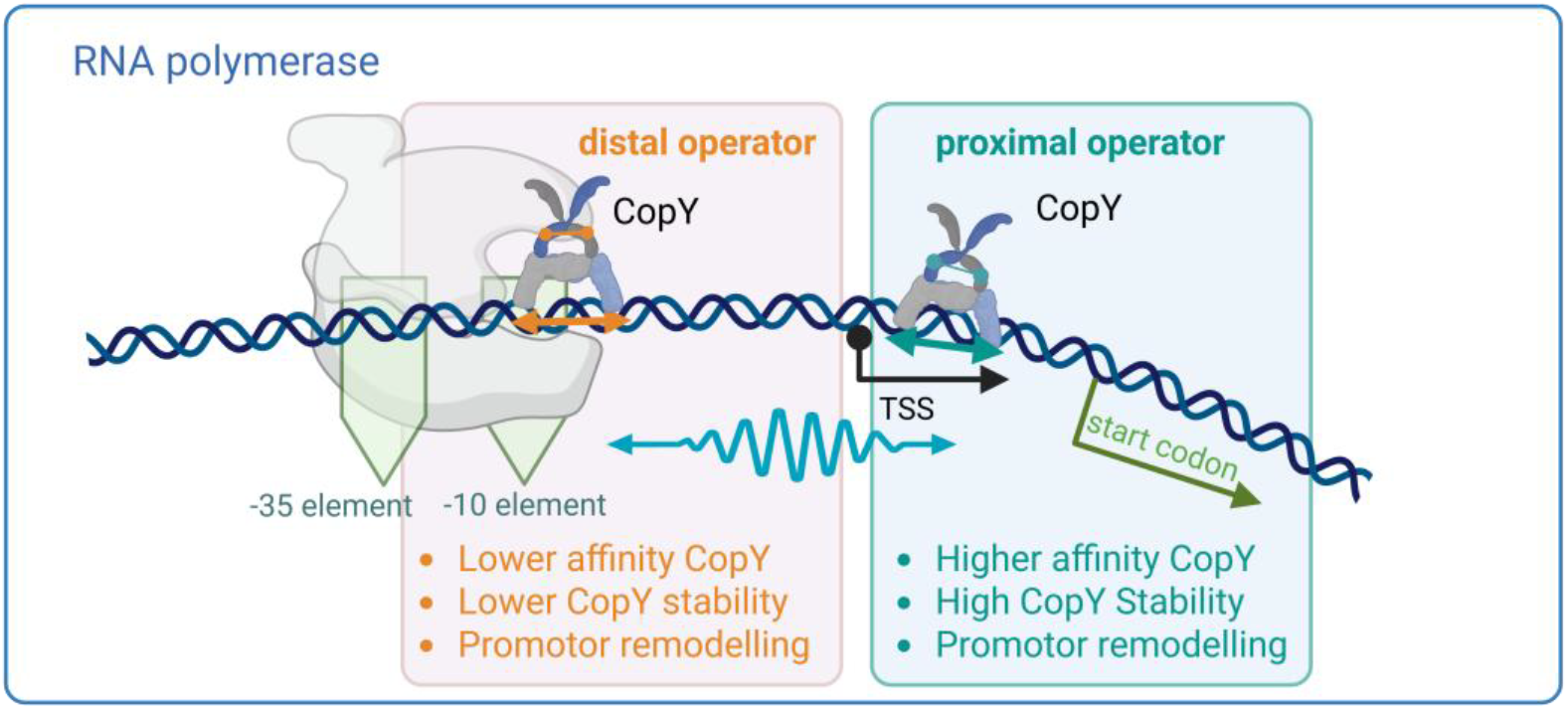

## Introduction

The transcriptional repressor CopY found in *Enterococcus* and *Streptococcus* exhibits distinctive mechanistic properties within the broader and structurally diverse class of metalloregulators (1-4). Unlike canonical transcription factors, which typically engage operator DNA as single dimers, CopY functions through simultaneous occupancy of two operator sites by two DNA-bound dimers (1,2,5,6). This dual-dimer configuration departs from the classical one-site repression paradigm and introduces an additional layer of regulatory complexity in bacterial copper homeostasis (7,8). It has been shown that two operator sites within the *cop* operon function in concert to mediate transcriptional repression (9,10). Two CopY dimers bind these operator sequences in the absence and presence of Zn(II), thereby maintaining the operon in a repressed configuration (5,11,12). Upon elevation of intracellular copper levels, Cu(I) acts as an allosteric modulator by displacing coordinated Zn(II), triggering conformational changes that reduce DNA-binding affinity and promote dissociation of CopY from both operator sites (5,6,13-16).

The *cop* operon promoter is composed of two discrete operator elements, a proximal and a distal site, each comprising a 16-26 base pair sequence that supports binding of CopY (Figure 1) (17). Notably, the distal operator overlaps with the canonical -35 and -10 promoter elements, whereas the transcriptional start site (TSS) is positioned between the two repressor binding sites, suggesting a spatial arrangement that enables direct interference with RNA polymerase recruitment and transcription initiation (Figure 1) (5). Despite the close proximity of these operator sequences, previous studies have reported an absence of measurable binding cooperativity, indicating that simultaneous occupancy does not arise from classical cooperative interactions (10). Importantly, the absence of thermodynamic cooperativity does not preclude functional or kinetic coupling between the two DNA-bound dimers. Rather, these observations suggest that transcriptional regulation is governed by intrinsic differences in binding affinity between CopY and the respective operator DNA sequences, resulting in site-specific contributions to repression. Quantitative analyses of CopY-DNA interactions support this model, revealing organism-specific differences in affinity between proximal and distal operator sites. In *Enterococcus hirae*, CopY binds more tightly to the proximal operator (K_D_ ≈ 2 nM) than to the distal site (K_D_ ≈ 5 nM) (5,6). A similar trend is observed in *Streptococcus pneumoniae*, where K_D_ values of ∼25 nM (proximal) and ∼55 nM (distal) have been reported (10), whereas *Lactococcus lactis* exhibits comparable affinities for both sites (K_D_ ≈ 32 nM) (18). Collectively, these findings indicate that differential site occupancy is primarily dictated by intrinsic binding affinities rather than cooperative interactions.

**Figure 1:**
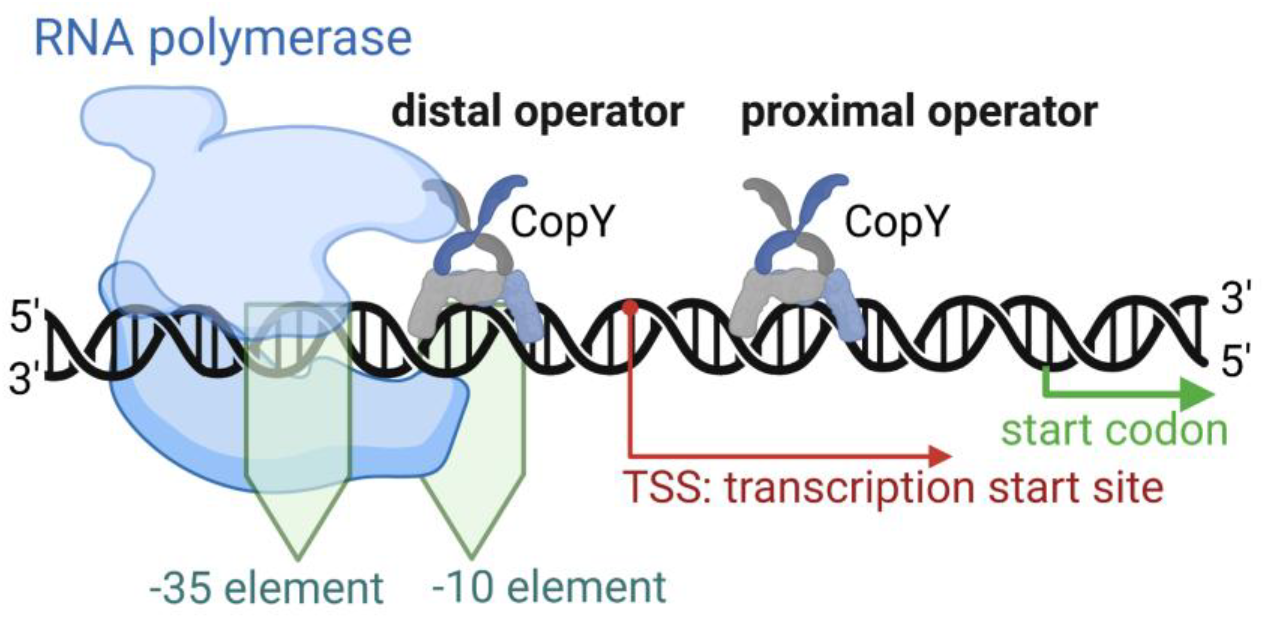
Schematic representation of the *cop* operon promoter region illustrating the spatial arrangement of distal and proximal operator sites and their interaction with CopY. The distal operator overlaps with the canonical −35 and −10 promoter elements, whereas the TSS is positioned between the two operator sequences. CopY binds DNA as two independent dimers, each occupying one operator site.

Despite the prevailing interpretation that CopY binding to dual operator sites occurs in a non-cooperative manner, the functional necessity of a dual-dimer architecture remains unresolved. The presence of two closely spaced operator sites, each independently occupied, suggests that regulatory coupling may not be adequately captured by classical thermodynamic definitions of cooperativity. Transcriptional regulation is governed not only by binding affinities but by the conformational states and flexibility of both partners, properties that are largely invisible to methods reporting on occupancy alone and difficult to capture by crystallography or cryo-EM, which favor well-ordered states. Electron paramagnetic resonance (EPR) distance measurements are uniquely suited to this problem, because spin labels placed at defined positions report distance distributions rather than single values, and therefore resolve conformational heterogeneity and its modulation in both protein and DNA, under solution conditions and without a size limitation (19-22). To gain mechanistic insight into CopY-mediated repression, we therefore employed various biochemical experiments coupled with EPR spectroscopy to monitor conformational and dynamical changes in both the CopY protein and its operator DNA.

## Results

Initially, we investigated the binding affinity of *S. pneumoniae* CopY for a long DNA fragment containing both distal and proximal binding sites, as well as for each site individually. We performed electrophoretic mobility shift assays (EMSAs) using three DNA constructs: the long DNA (including both distal and proximal sites), and two shorter sequences containing either the distal site DNA, or the proximal site DNA. Figure 2A presents the EMSA analysis of CopY binding to the long DNA as a function of increasing CopY concentration. The appearance of multiple shifted bands indicates that the long DNA fragment contains two independent CopY-binding sites that can be sequentially occupied by CopY dimers.

**Figure 2:**
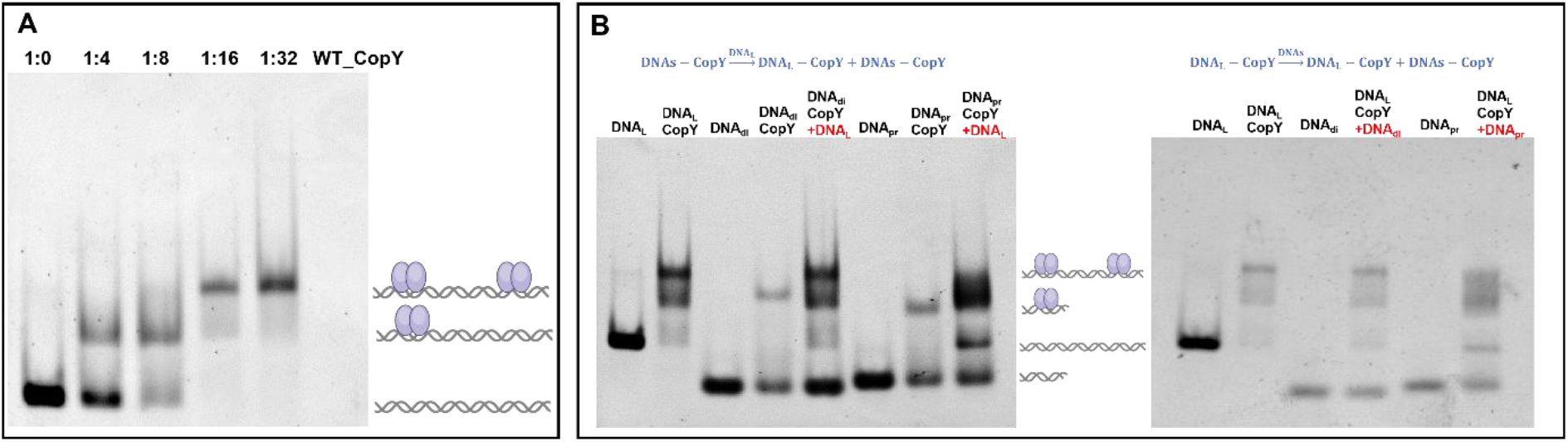
EMSA gel of **(A)** binding of CopY to a long DNA fragment as a function of increasing CopY concentration. The appearance of multiple shifted species indicates sequential occupancy of two distinct binding sites by CopY dimers. **(B)** Competition EMSA experiments. **Left panel:** CopY was first bound to either the distal or the proximal short DNA fragment, and the long DNA was subsequently added. **Right panel:** CopY was first incubated with the long DNA fragment, followed by addition of either the distal or the proximal short DNA fragment.

To compare the relative affinities of CopY for the distal and proximal sites, competition experiments were performed. First, CopY was incubated with either the distal or proximal short DNA fragment (Figure 2B, left panel), followed by the addition of the long DNA. The EMSA results showed that the long DNA displaced CopY from the distal DNA more efficiently than from the proximal DNA. This observation suggests that CopY binds more strongly to the proximal site than to the distal site. In a complementary experiment, CopY was first incubated with the long DNA (Figure 2B, right panel), subsequently the distal or proximal short DNA fragment was added. Consistent with the previous results, the proximal DNA competed more effectively with the long DNA for CopY binding than did the distal DNA. Together, these findings indicate that CopY exhibits a higher binding affinity for the proximal site compared to the distal site.

To further explore the stability of the CopY-DNA complex, circular dichroism (CD) experiments were conducted as a function of temperature for wild-type (WT) CopY alone and in complex with each of the three DNA constructs: long DNA, distal DNA, and proximal DNA (Figure S1, SI). Figure 3A and Figure 3B show the CD spectra recorded at different temperatures for CopY alone and the CopY-long DNA complex, respectively. The CD spectrum of CopY is characteristic of a largely α-helical structure, exhibiting negative peaks at 210 and 220 nm. As expected, the intensity of these negative peaks decreases upon heating. The temperature-dependent CD data were analyzed to compare the apparent thermal stability of CopY alone and in complex with the long, distal, or proximal DNA constructs (see SI). At each temperature, the CD signal was averaged over 220-224 nm to reduce single-wavelength noise. A common temperature range of 20-85°C was used for all samples. Data collected at 90°C were excluded because of the anomalously large change observed between 85 and 90°C. The data were analyzed using two complementary methods. First, a model-free analysis was performed in which each trace was normalized using its measured values at 20 and 85°C. The observed midpoint temperature, T_50_, was defined as the temperature at which the observed fraction is equal to 0.5 and was determined by linear interpolation between the two surrounding experimental points (Figure 3C). In the second analysis, the four non-normalized traces were fitted simultaneously (Figure 3D). The van’t Hoff relationship was used to describe the temperature-dependent equilibrium between the native and unfolded states (23), with a common van’t Hoff enthalpy shared among all four datasets (see SI). Both analyses indicate that DNA binding increases the apparent thermal stability of CopY (Figure 3, table). The model-free midpoints were 51 °C for CopY alone, 62 °C with distal DNA, 66 °C with proximal DNA and 66 °C with long DNA; the global fit gave 51 ± 1, 64 ± 1, 72 ± 2 and 73 ± 3 °C, respectively. The two analyses yield the same rank order. Proximal and long DNA stabilize CopY to a comparable extent, whereas distal DNA produces less stabilization than proximal DNA.

**Figure 3:**
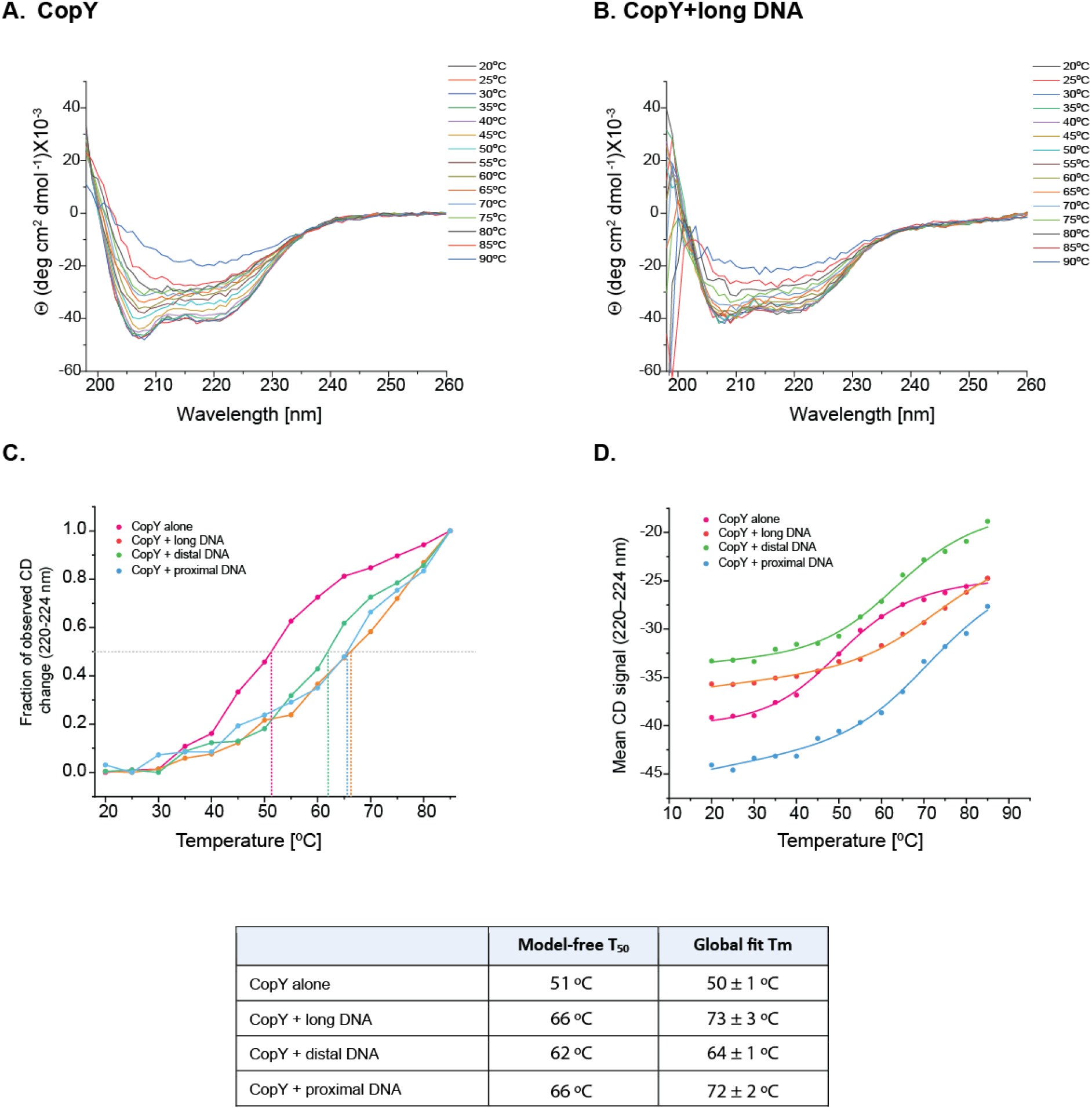
CD spectra of (**A**) CopY and (**B**) the CopY-long DNA complex recorded at different temperatures. (**C**) Change in ellipticity of the mean value between 220-224 nm as a function of temperature (20-85 °C) for all four samples. (**D**) Global fit of the thermal denaturation curves sharing a single van’t Hoff enthalpy (ΔH = 107 ± 17 kJ mol^−1^). The table at the bottom lists the model free T_50_ and Tm values obtained from the fit for each sample.

To further elucidate the repression mechanism of CopY, we employed an EPR-based DNA spin-labeling strategy that enables the detection of subtle DNA conformational changes associated with protein binding (24,25). Four DNA constructs derived from the isolated proximal and distal operator sequences were designed, each containing a single CopY-binding site and therefore capable of binding only one CopY dimer (Figure 4, top panels, and Figure S2, SI). The spin-label positions were strategically selected to probe distances across the CopY-binding site, more specifically one was positioned within the center (Fig. 4 purple star, *Cop_10c/9c*), and the second one was positioned flanking the binding site (Fig. 4 blue star, *Cop_14p*). This design allowed us to distinguish between local structural perturbations within the operator region and more global DNA conformational changes induced by CopY binding. EMSA experiments confirmed binding of WT-CopY to the spin-labeled DNA (Figure S3, SI).

**Figure 4:**
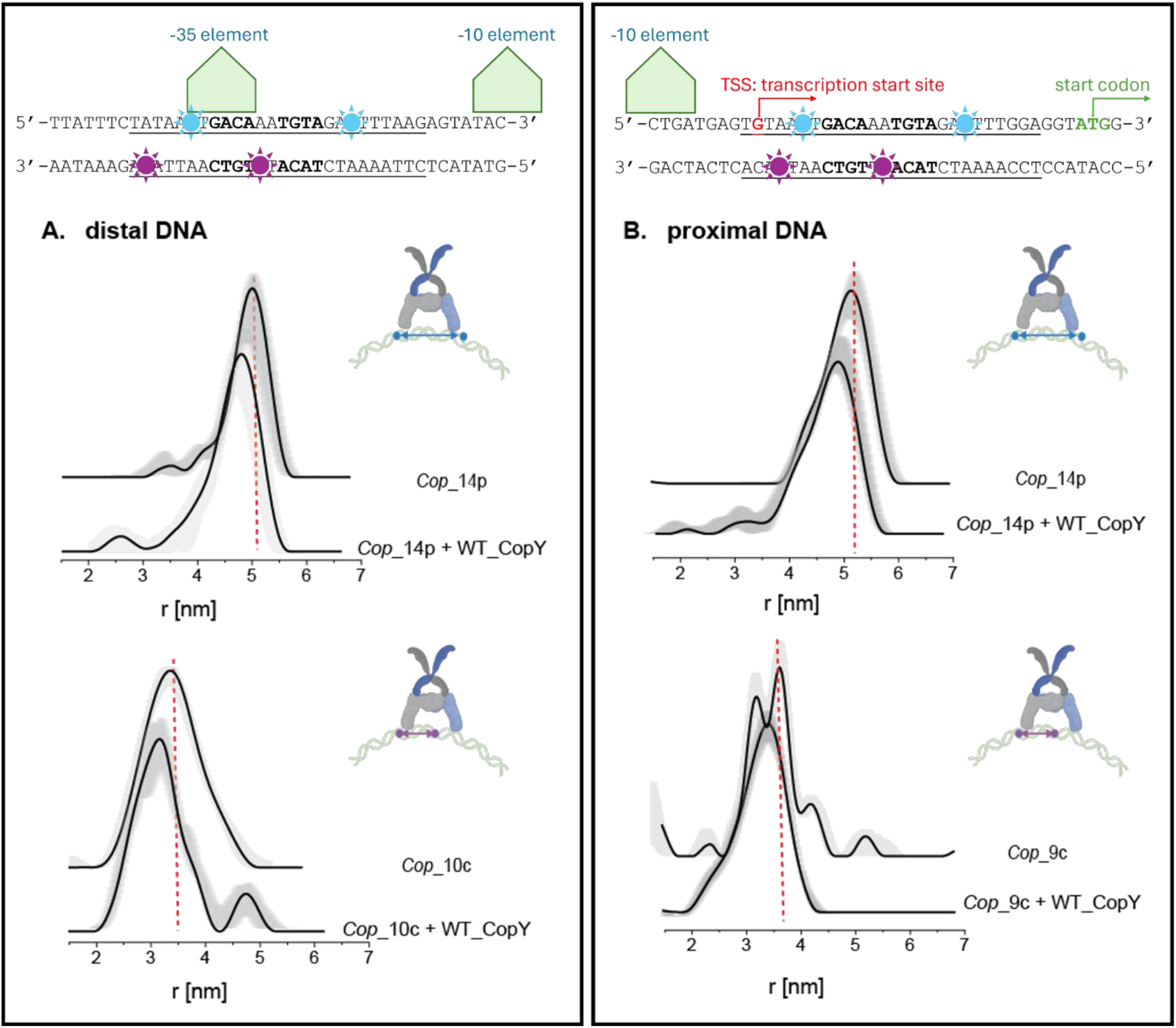
**Top:** Schematic representation of the double-stranded DNA sequences used to probe the structural dynamics of the *cop* regulatory region via EPR spectroscopy. Blue stars denote labeling sites positioned in the flanking regions outside the core CopY binding motifs. Purple stars indicate labeling sites located within the center of the operator core. These sites are symmetrically coordinated relative to the conserved inverted repeat motifs (GACAA and TGTAGA), which facilitate independent homodimer binding of CopY. The spatial overlay with the -35 and -10 promoter elements and the TSS illustrates the topography through which CopY-induced DNA deformation orchestrates transcriptional repression. DEER distance distribution for (**A**) distal DNA and (**B**) proximal DNA in the absence and presence of WT CopY. Red lines were drawn to guide the eye.

Using site-directed spin labeling in combination with Double electron-electron resonance (DEER) spectroscopy, we observed that CopY binding induces conformational rearrangements in both the distal (Figure 4A, Figure S4-5, SI) and proximal (Figure 4B, Figure S6-7, SI) operator sequences. The most pronounced effects were detected for DNA constructs in which both spin labels were positioned outside the CopY-binding site (*Cop_14p*). In these constructs, CopY binding resulted in a reduction of approximately 0.4-0.6 nm in the average inter-spin distance for both the proximal and distal operators. These distance changes indicate that CopY binding promotes DNA compaction or bending at both operator sites. Some minor changes were detected upon protein coordination in the room-temperature (RT) continuous-wave EPR (CW-EPR) spectra (Figure S8, SI). Notably, although CopY binds the two operators with different affinities and they confer different thermal stabilization, the DNA itself undergoes a similar structural rearrangement at both sites, suggesting that DNA deformation is a common feature of CopY-mediated repression.

To investigate the conformational and dynamic changes that occur in CopY upon binding to different DNA fragments, we employed site-directed spin labeling coupled with EPR spectroscopy. CopY contains three native cysteine residues (C52, C86, and C101). Previous studies identified C101 as the residue most sensitive to DNA binding (13). Therefore, we generated the double mutant CopY_C52A_C86A, leaving C101 as the sole labeling site for attachment of the nitroxide methanethiosulfonate spin-label (MTSSL). CD and EMSA confirmed that the mutant preserved the secondary structure and the binding to the DNA (Figure S9, SI). DEER experiments were performed on spin-labeled CopY_C52A_C86A in the absence and presence of long, distal, and proximal DNA fragments. The DEER time-domain traces are shown in Figure 5A,B. For the apo protein, the DEER signal exhibits only weak dipolar modulation, indicative of high conformational flexibility, and a broad distribution of interspin distances. Binding to the long or proximal DNA results in a slightly more pronounced dipolar modulation, suggesting a small reduction in conformational heterogeneity, whereas binding to the distal DNA produced no detectable change.

**Figure 5:**
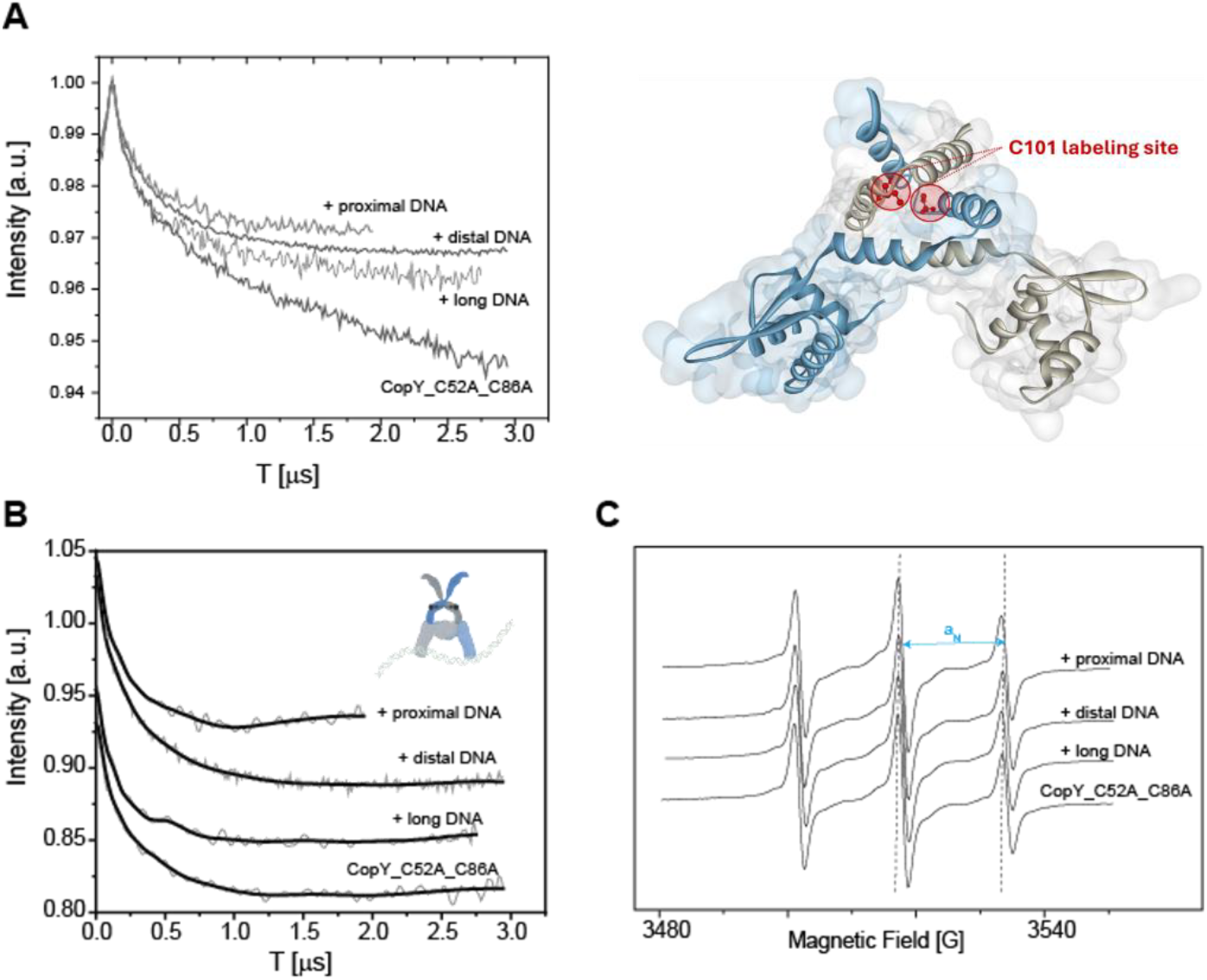
Q-band DEER time-domain traces before (**A**) and after background subtraction (**B**) of MTSSL-labeled CopY_C52A_C86A in the absence of DNA and in the presence of long, distal, or proximal DNA fragments. **(C)** X-band CW-EPR spectra of MTSSL-labeled CopY_C52A_C86A recorded at room temperature. Right of (A): model of CopY with a truncated C-terminus bound to DNA. CopY model was generated with Swiss-Model Server and is based on the structure of BlaI repressor from *Staphylococcus aureus* (PDB-ID: 1SD7), the labeling site C101 is highlighted in red (26,27).

To further characterize local changes around the spin-label site, RT CW-EPR spectra were recorded (Figure 5C). The spectra indicate that the nitroxide label attached to C101 remains highly mobile under all conditions, suggesting that the local flexibility of the labeling site is largely preserved upon DNA binding. Nevertheless, a slight decrease in the hyperfine coupling constant (a_N_) was observed upon binding to the long or proximal DNA fragments. This change is consistent with the spin label experiencing a more hydrophobic environment, which may reflect a subtle change in the local environment of C101.

The pronounced flexibility of the CopY metal binding domain was confirmed by another labeling approach, using the rigid dHis-Cu(II)-NTA label in the CopY_A95H_V99H variant. Because this label is held bipedally against the protein backbone, it eliminates the rotameric freedom of the nitroxide tether and typically yields substantially narrower distance distributions, so any breadth that remains reports on the protein rather than on the label (28-35). Figure 6 shows the time domain Relaxation-Induced Dipolar Modulation Enhancement (RIDME) traces and the measured distance distribution. We chose RIDME over DEER for these measurements to achieve higher modulation depths. The distributions are all complex and display a large range of distances although there appears to be a small decrease in the range of measured distances when the protein is complexed to the long DNA and proximal DNA (the distance ranges go from ca. 2.2 – 4.9 nm to 2.2 – 4.4 nm).

**Figure 6:**
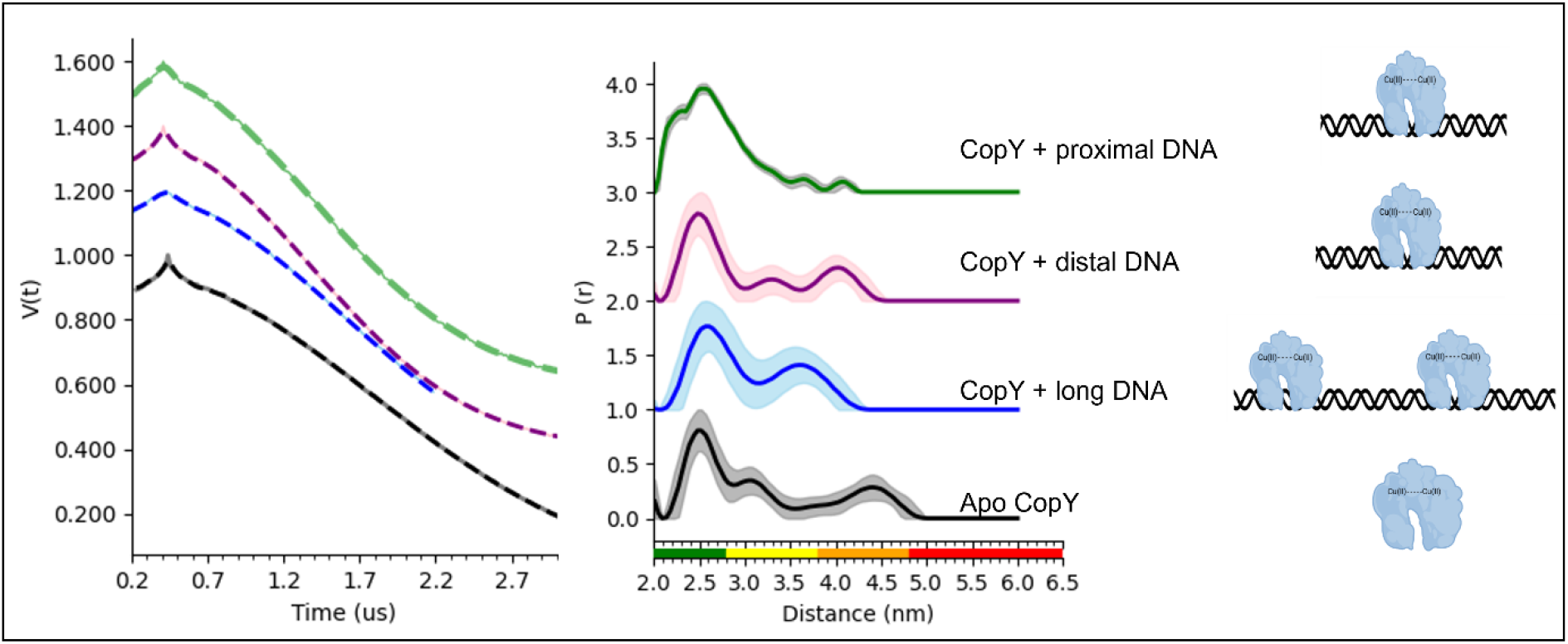
Q-band RIDME time-domain of Cu(II)-NTA bound to CopY_A95H_V99H mutant in the absence and in the presence of long, distal, or proximal DNA fragments, and the corresponding distance distribution functions. The data was analyzed using the DeerAnalysis2026 program using the DeerLab framework. Generalized cross-validation was chosen as the regularization parameter for all data analysis. The color bar indicates reliability ranges (green: shape reliable; yellow: mean and width reliable; orange: mean reliable; red: no quantification possible).

Overall, the distance measurements using both the MTSSL and the dHis-Cu(II) labels indicate that the metal binding domain in the two subunits of the CopY protein exhibit a large conformational space with respect to each other. In addition, the CW-EPR data on MTSSL shows that the backbone reorients on a rapid timescale. Together these results are intriguing for this metalloregulator which is nominally bound to the DNA holding the DNA in the repressed state. We hypothesize that the large conformational range and fast dynamics confers advantages in the efficient search for free Cu(I) for this regulator. Notably prior work on an activator CueR has shown that the metal binding domain is much more rigid (36). These results highlight the differences between two similar copper transcription factors that operate distinctly – e.g. CopY is a transcriptional repressor while CueR is an activator. A working model for the promoter states is summarized in Figure 7.

**Figure 7:**
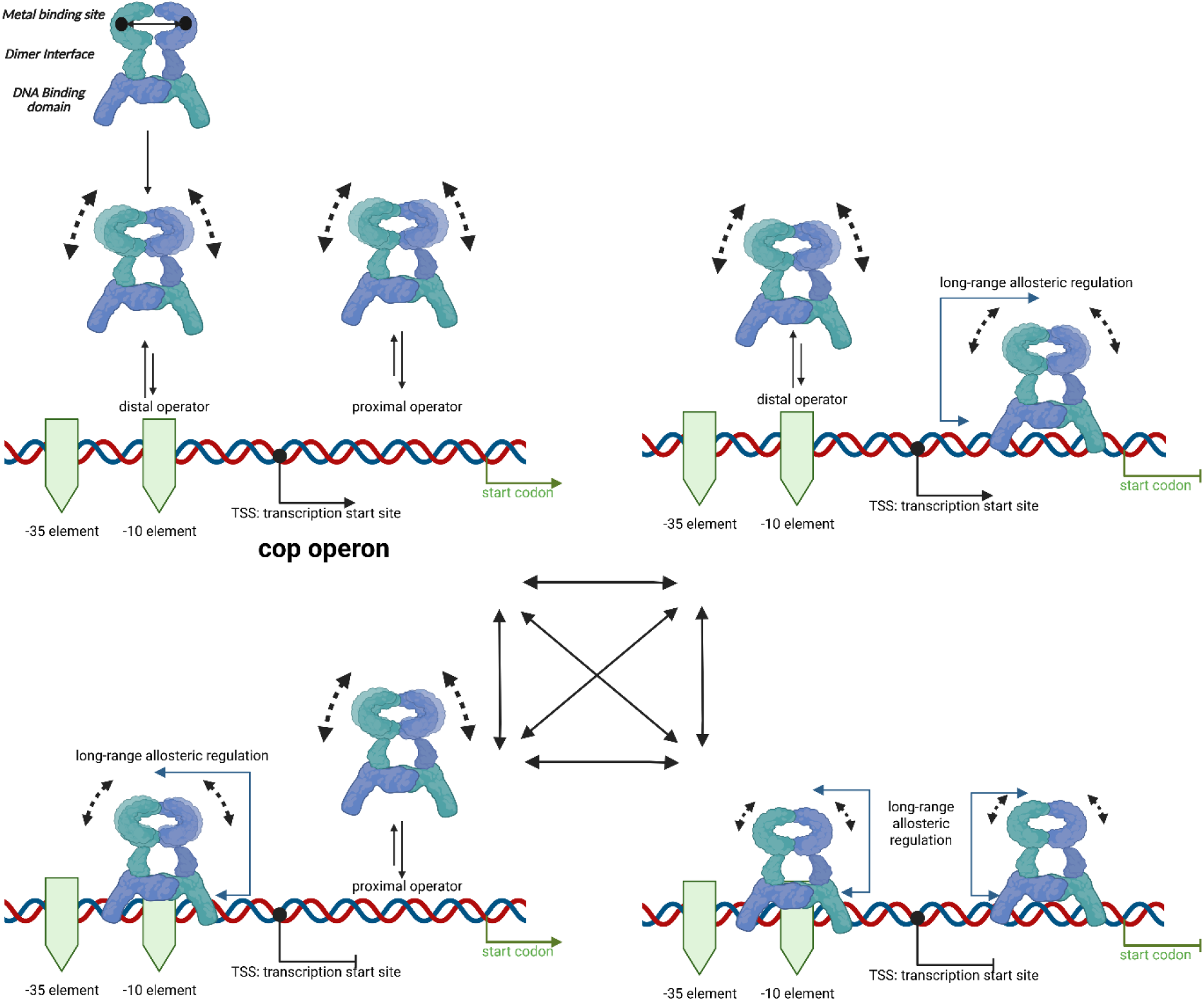
Model for CopY-mediated repression of the *cop* operon, represented as an equilibrium between four states of the promoter. **Top left**: in the unbound state, apo CopY dimers are free in solution; the dashed arrows indicate the wide range of geometries sampled by the metal-binding domain. **Top right and bottom left:** occupancy of the proximal operator alone and of the distal operator alone. Because the distal operator overlaps the −35 and −10 elements, its occupancy interferes directly with RNA polymerase, whereas the proximal operator provides the higher-affinity anchor. **Bottom right**: the fully repressed state, in which both operators are occupied. In all DNA-bound states, the metal-binding domain remains highly flexible, with only marginal narrowing of its conformational range relative to the free protein. The double-headed arrows in the center indicate that the four states interconvert rather than following an obligatory order of binding. Schemes were created using BioRender.

## Discussion

The present study provides new mechanistic insights into how CopY regulates copper homeostasis through two non-equivalent operator sites within the *cop* promoter. While previous studies established that two CopY dimers sequentially occupy the distal and proximal operators, the structural consequences of these interactions remained unclear (6). By combining EMSA, CD spectroscopy, protein spin-labeling, and DNA-based EPR measurements, we demonstrate that the two operator sites play distinct but complementary roles in repression, utilizing the DNA scaffold as a structural conduit (37,38).

The EMSA competition experiments confirm that CopY binds the proximal operator with higher affinity than the distal operator. This observation is consistent with previous reports for CopY homologs and suggests that the two operator sites are not functionally equivalent (5). Importantly, the higher affinity of the proximal site is mirrored by the CD measurements, which show greater thermal stabilization of CopY when bound to the proximal DNA compared with the distal DNA. The increased unfolding temperature observed in the presence of proximal DNA indicates that this operator stabilizes the CopY structure more effectively, suggesting the formation of a more favorable protein-DNA interface.

A particularly intriguing outcome of this work is the apparent decoupling between DNA-binding affinity and DNA deformation. Although CopY binds the distal operator with lower affinity, DNA-based DEER measurements reveal that both distal and proximal DNA operators undergo comparable structural rearrangements upon protein binding. These findings indicate that DNA deformation is not dictated solely by binding strength; rather, DNA bending or compaction appears to be an intrinsic consequence of operator recognition by CopY. The capacity of the distal operator to undergo structural remodeling despite its weaker affinity suggests that DNA deformation represents a primary functional output of repression, rather than a passive byproduct of tight binding.

The observation that CopY induces localized DNA compaction distinguishes it from other well-characterized metalloregulators, particularly members of the MerR family such as CueR (25,34,39-41). MerR-family regulators utilize DNA distortion to activate transcription by realigning promoter elements that are sub optimally spaced for RNA polymerase recognition (42-44). In contrast, the structural rearrangements observed here appear to serve a repressive function. Because the distal operator directly overlaps the -35 and -10 promoter elements, CopY-induced distortion of this region likely interferes with RNA polymerase binding and/or promoter melting. Thus, although both CopY and MerR-family proteins exploit DNA deformation as a regulatory mechanism, the functional consequences of the deformation are fundamentally different.

Taken together, the results support a model in which the two operator sites perform specialized roles within the repression complex. The proximal operator primarily contributes to stable CopY binding and thermal stabilization, whereas the distal operator, which overlaps the −35 and −10 elements, contributes to promoter remodeling and transcriptional interference. This division of labor provides a mechanistic explanation for why two independently occupied operators are maintained despite the apparent absence of classical thermodynamic cooperativity.

Figure 7 outlines this proposed working model as an equilibrium among four states of the *cop* promoter. In all four states, the metal-binding domain of CopY samples a broad conformational ensemble, and DNA binding produces only marginal changes in this ensemble. The singly bound states are non-equivalent: the higher-affinity proximal site acts as the primary anchor, whereas the distal operator overlaps the promoter elements, where repressor binding directly interferes with RNA polymerase recruitment. Whether occupancy of one operator influences the dimer bound at the adjacent site cannot be resolved by the present data and remains to be tested. Importantly, the four states are expected to interconvert rather than follow an obligatory binding order, so that the relative populations of these states, rather than a fixed sequence of events, set the depth of repression. The persistent flexibility of the metal-binding domain in the DNA-bound states may, in turn, keep the repressor poised to respond rapidly to Cu(I).

Such functional asymmetry may provide important physiological advantages. Complete repression of copper-export genes under normal growth conditions minimizes unnecessary energy expenditure. However, maintaining a promoter architecture that is capable of rapid derepression upon copper stress is equally important. The weaker distal operator may therefore function as a dynamic regulatory element that permits transient fluctuations in occupancy while the proximal operator remains stably bound. Such an arrangement would allow low levels of basal transcription and would enable rapid activation of copper detoxification pathways following a sudden increase in intracellular copper.

Collectively, these observations suggest that CopY operates as a structurally dynamic regulatory system in which differential operator recognition, protein conformational stabilization, and DNA remodeling are integrated to control transcription. The proximal and distal operators thus act as complementary regulatory modules that together generate efficient and responsive control of copper homeostasis.

## Conclusions

This work provides a structural framework for understanding how CopY regulates transcription through dual operator recognition. Biochemical experiments demonstrate that CopY binds the proximal operator with higher affinity than the distal operator, while CD measurements reveal that proximal DNA binding confers greater thermal stabilization of the protein. Despite these differences in protein response, DNA-based DEER measurements reveal that both operator sites undergo comparable structural rearrangements upon CopY binding. These findings indicate that DNA deformation and binding affinity are not directly related. Moreover, EPR measurements show that the metal-binding domain of CopY is highly flexible and conformationally heterogeneous, with only marginal changes upon DNA binding, suggesting that the DNA-bound repressor keeps its metal site dynamic and accessible to Cu(I). Here, the proximal operator appears to serve primarily as a high-affinity anchoring site that stabilizes the repressor complex, whereas the distal operator, which overlaps the core promoter elements, contributes directly to transcriptional repression. This mechanism expands the current understanding of bacterial metalloregulation and highlights how promoter architecture can integrate protein dynamics and DNA remodeling to achieve robust control of metal homeostasis.

## Materials and Methods

### CopY cloning, expression, and purification

The *copY* gene was amplified from *S. pneumoniae* genomic DNA by PCR using primers containing specific *copY* sequences and flanking regions corresponding to the pET28a vector sequence. The forward primer used was 5′-GCGGGTGAAAACCTGTACTTCCAGGGTATGCAGATTTCAGATGCAGAATGG-3′, and the reverse primer was 5′-GCTTTGTTTAGCAGCCTAGGTATTAATCAATTACATACAATTACATCTTACTTCTGT-3′.

The amplicon was cloned into the pET28a vector using restriction-free cloning (45). pET28a is a cloning and expression vector that allows overexpression of CopY fused with maltose-binding protein (MBP) and a His-tag. The engineered sequence was expressed in *Escherichia coli* BL21 cells grown to an optical density of 0.8 at 600 nm and induced with 1 mM isopropyl-β-thiogalactopyranoside (IPTG) at 20 °C overnight with shaking. The cells were harvested by centrifugation, and the pellet was resuspended with a buffer containing 25 mM Tris-HCl, pH 7.4, 250 mM NaCl, 10 mM imidazole, and 1% Triton X-100. The *E. coli* cells were sonicated (10 min total, 10 sec pulses, 35% amplitude) and harvested by centrifugation at 14,000 rpm for 40 min. The soluble fraction of the CopY-containing lysate was purified by Ni-NTA agarose beads according to the manufacturer’s protocol. The fusion protein was eluted with buffer containing 25 mM Tris-HCl, pH 7.4, 250 mM NaCl, and 300 mM imidazole. The eluted fraction was then incubated and dialyzed overnight with tobacco etch virus protease (TEV) and 5 mM DTT to cleave the fusion protein and remove the His-tag. Finally, purified CopY fractions were collected from a second Ni-NTA agarose bead column using a buffer containing 25 mM Tris-HCl, pH 7.4 and 250 mM NaCl. Each purification step was verified by SDS-PAGE and Coomassie blue staining. Expression and purification of CopY_C52A_C86A was carried out using the same protocol. The expression and purification of the dHis mutant of CopY were performed with the modification that 25 mM HEPES, 250 mM NaCl, pH 7.4 were used as the purification buffer to enable subsequent spin labeling without an additional buffer exchange step.

### CopY protein spin-labeling

CopY was first incubated overnight with 10 mM DTT and Cu(I) (tetrakis (acetonitrile) copper(I) hexafluorophosphate). DTT was dialyzed out using a GeBaFlex-tube with a molecular weight cut-off of 1 kDa against protein buffer at 4°C overnight. Next, Cu(I) and an excess of (1-oxyl-2,2,5,5-tetramethylpyrroline-3-methyl) methanethiosulfonate (MTSSL, TRC) were added to the protein and the solution was then vortexed overnight at 4 °C. The free spin-label and Cu(I) ions were removed by several dialysis cycles over 4-5 days. For Cu(II)-NTA labeling, 80 μM dHis CopY monomer was mixed with an equimolar concentration of Cu(II)-NTA (80 μM) in a 500 μL microcentrifuge tube and incubated at 4°C for 30 min, using the protocol described in detail earlier (46). For EPR measurements, spin-labelled proteins were diluted in protein buffer with 20% glycerol.

### DNA spin-labeling

High-performance liquid chromatography (HPLC)-purified double strands DNA (Table 1) with a modified thymine base was purchased from IDT (Syntezza Bioscience, Jerusalem, Israel) and dissolved in double deionized water, resulting in a 100 μM stock concentration. For DNA precipitation, 2 nmol DNA was mixed with 2 μL 3M NaOAc (pH 5.2). Subsequently, 60μL ice-cold Et-OH was added and incubated for 1 h at 20°C. Upon centrifugation at 14,000 rpm and 4°C for 25 min, a white pellet confirmed the precipitation of DNA. The supernatant was carefully removed without disturbing the pellet. The Eppendorf tube containing the pellet was dried overnight by lyophilization. For the labeling reaction, 10 μL of labeling buffer (0.1 M sodium tetra borate, pH 8.5) was added. Next, nitroxide-NHS-ester was added and mixed gently by pipetting 10 μL of 8 mM solution in dry Dimethylsulfoxide (DMSO). The reaction was incubated at 85°C for 1 h while shaking at 500 rpm. The reaction was quenched by the addition of 1.5 μL 3 M NaCl and 50 μL ice-cold EtOH (stored at -20°C). Then, the DNA mixture was precipitated by EtOH precipitation at 20°C for 1 h. The solution obtained was centrifuged at 14,000 rpm and 4°C for 25 min. After centrifugation, the supernatant was carefully removed, and the resulting pellet was dried by lyophilization overnight. For final purification, the pellet was dissolved in 20 μL Triethylammonium acetate (TEAA) buffer and subjected to HPLC purification by reversed-phase chromatography using a XBridge BEH C18 column, 130 Å, 2.5 μm, 4.6 × 50 mm column (Waters, Milford, MA, USA). The sample (20 μL) was loaded onto a C18 column using HPLC (Agilent 1200 System). DNA strands were eluted by the following program: isocratic flow 95% buffer A (TEAA 0.1 M, pH 6.5) and 5% buffer B (100% acetonitrile) for 7.5 min; gradient flow to 80% buffer A and 20% buffer B for 7.5 min; gradient flow from 80% to 0% buffer A and to 100% buffer B for 5 min; isocratic flow of 100% buffer B for 2 min; and gradient flow to 95% buffer A and 5% buffer B in 3 min followed by isocratic flow of 95% buffer A and 5% buffer B for 2 min. Separation was achieved at a flow rate of 1 mL/min. The fractions with labeled DNA were pooled and dried by lyophilization overnight. The samples were subjected to LC-tandem mass spectrometry to confirm successful labeling of the DNA strands. The labeled DNA double strands (Table 1) were stored at -20°C with 20% glycerol until further use.

**Table 1:**
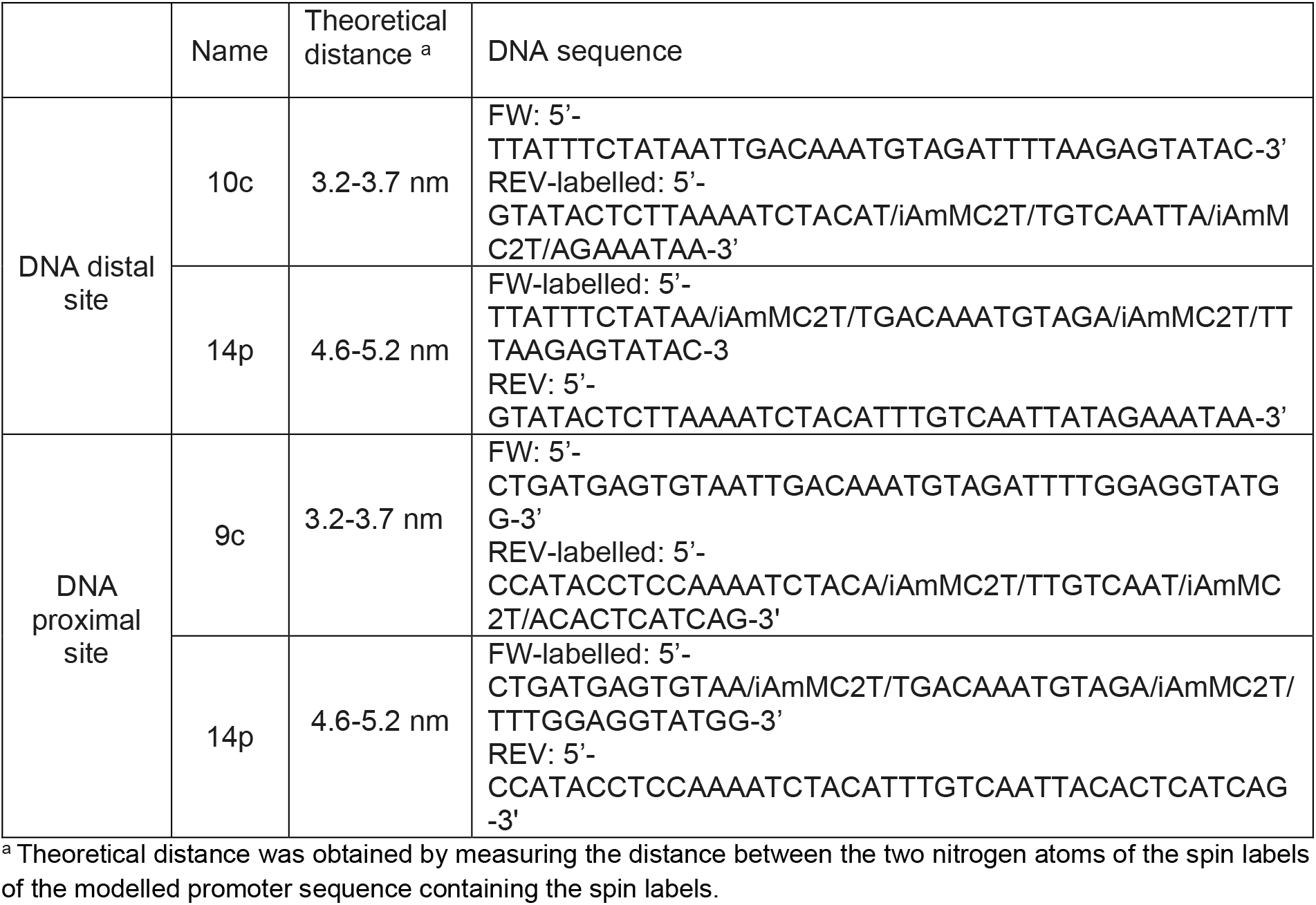
Spin-labeled DNA sequences used in this study.

### Electrophoresis Mobility Shift Assay (EMSA)

Electrophoretic mobility shift assays (EMSA) were performed using 5% native acrylamide gels. Wild type CopY protein (WT_CopY) was incubated with *cop* operon DNA in the presence of 20% glycerol at room temperature for 30 min. For competition assays, WT_CopY was pre-incubated with one DNA fragment (long, distal, or proximal) for 30 min, followed by addition of a second fragment and further incubation under the same conditions. Complexes were resolved in TAE buffer (40 mM Tris-HCl, pH 8, 20 mM acetic acid, 1 mM EDTA at 4 °C and 80 V for 1 h. Gels were stained with 1 μg/mL ethidium bromide and imaged using a Gel Doc EZ System.

### CD

CD measurements were performed using a Chirascan spectrometer (Applied Photophysics, UK) at temperature range of 20-90 °C. Measurements were carried out in a 1 cm optical path length cell. The data were recorded from 190–260 nm with a step size and a bandwidth of 1 nm.

### CW-EPR

Continuous wave EPR (CW-EPR) experiments were performed on X-band E500 Elexsys Bruker spectrometer. The spectra were recorded at RT with microwave power of 20.0 mW, time constant of 60 ms, modulation amplitude of 1.0 G, and a receiver gain of 60 dB. The samples were measured in 0.8 mm capillary quartz tubes (vitrocom).

### DEER

The DEER experiment π/2(v_obs_) − τ1 − π(v_obs_) − t′ − π(v_pump_) − (τ1 + τ2 − t′) − π(v_obs_) − τ2 − echo was carried out at 50 ± 1.0 K on a Q-band Elexsys E580 spectrometer (equipped with a 2-mm probe head) and AmpQ 50W TWT. A two-step phase cycle was employed on the first pulse. The echo was measured as a function of t′, whereas τ2 was kept constant to eliminate relaxation effects. The durations of the observer π/2 and π pulses were 12 ns and 24 ns, respectively. The duration of the π pump pulse was 24 ns, and the dwell time was 16 ns. τ1 was set to 200 ns. The observer frequency was 33.8 GHz, the pump frequency was 33.86 GHz, and the magnetic field was 12025 G. The samples were measured in 1.6 mm capillary quartz tubes (Wilmad-LabGlass). The data was analyzed using the DeerAnalysis 2019 program (47,48).

### 5 -pulse RIDME and data analysis

All Q-band RIDME measurements were performed at 30K using a Bruker ElexSys E580 FT/CW spectrometer with a Bridge 12 Technologies QLP Q-band resonator, Oxford ITC503 temperature controller, Oxford CF935 dynamic continuous-flow cryostat, and Oxford LLT 650 low-loss transfer tube, a 300 W TWT amplifier, and a Bruker SpinJet AWG.

A five-pulse RIDME sequence (49) of π/2–τ_1_–π–(τ_1_+t)–π/2–T_mix_–π/2–(τ_2_-t)–π–τ_2_–echo with a π/2 pulse of 10 or 6ns and a corresponding π pulse of 20 or 12 ns. All experiments were performed at the maximum of the Cu(II)-NTA field-swept echo-detected spectrum, as described previously (50). During the RIDME experiments, t was incremented in 16 ns steps, yielding 188, 157, or 125 data points, depending on the dipolar evolution time. An 8-step phase cycle was employed, and ^2^H modulation was suppressed using 16-step τ_1_ averaging. The mixing time, T_mix_, was set to 0.7 * T_1_, and the shot repetition time was adjusted to 3-4 * T_1_.

All five-pulse RIDME data were analysed using DeerAnalysis2026, using the DeerLab framework (48,51). A stretched-exponential function was used to model the background contribution. The background dimensionality was not constrained during fitting; however, the optimized dimensionality was consistently between 5 and 6, in agreement with values previously reported for RIDME experiments (50,52). Generalized cross-validation was chosen as the regularization parameter for all data analysis.

## Acknowledgements

Schemes of protein and DNA were created using BioRender.

## Supplementary Data Statement

Additional experiments are available in the supporting information file.

## Author Contribution statement

MH, AB, and YS conducted the experiments and analyzed the data. YS also contributed to the mass spectrometry and CD experiments and their analysis. LH and LGA assisted with the study design and data interpretation. SR and SS conceived, designed, and supervised the research. All authors contributed to writing and revising the manuscript and participated in ongoing scientific discussions.

