## Supplementray Information for "CopY as a Transcriptional Duet Orchestrating Copper Homeostasis via Asymmetry-Dependent DNA Affinity"

**Supporting Information**

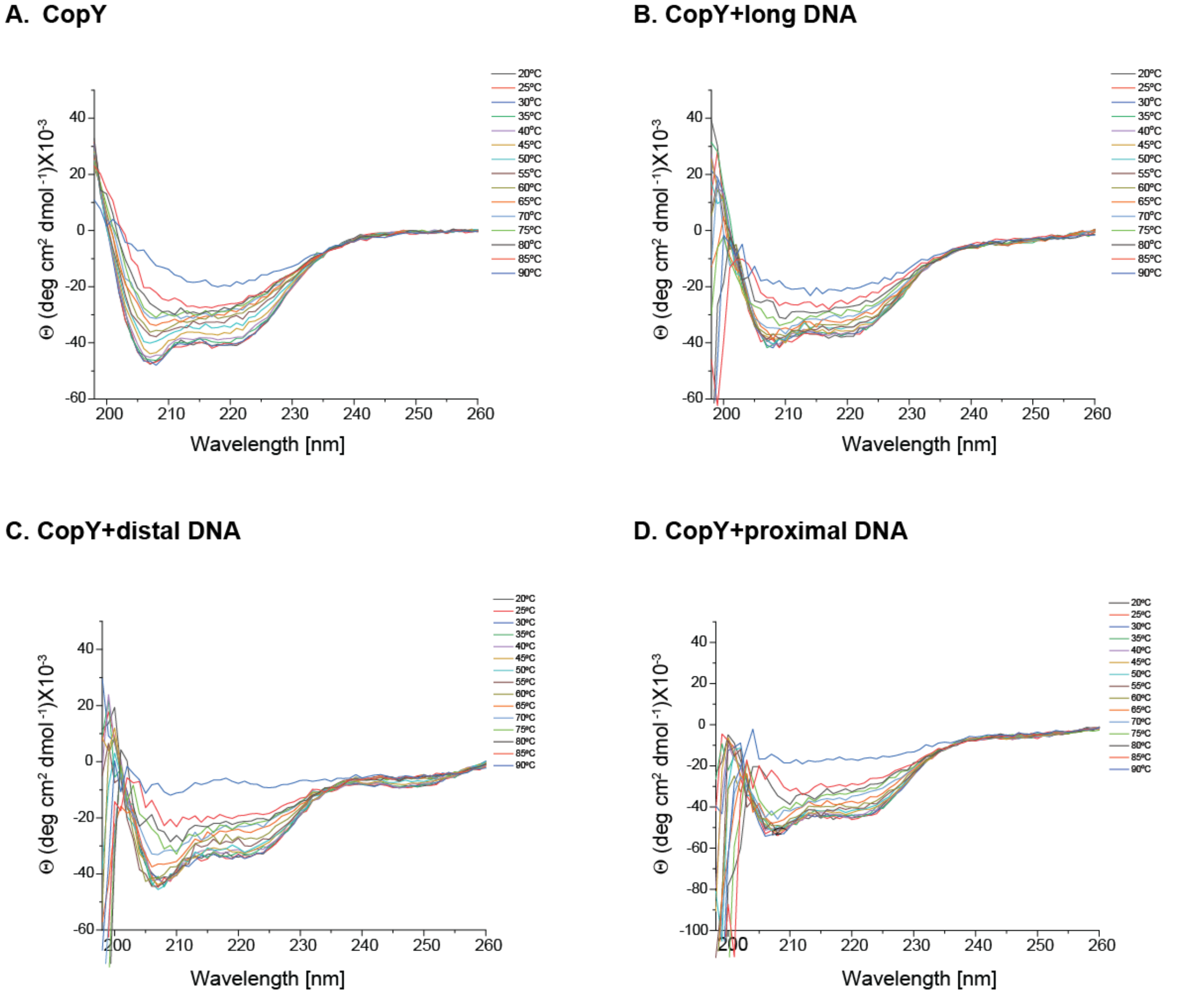


**Figure S1**: CD spectra recorded at different temperatures for (**A**) CopY, (**B**) CopY-long DNA complex, (**C**) CopY-distal DNA complex, and (**D**) the CopY-proximal DNA complex.

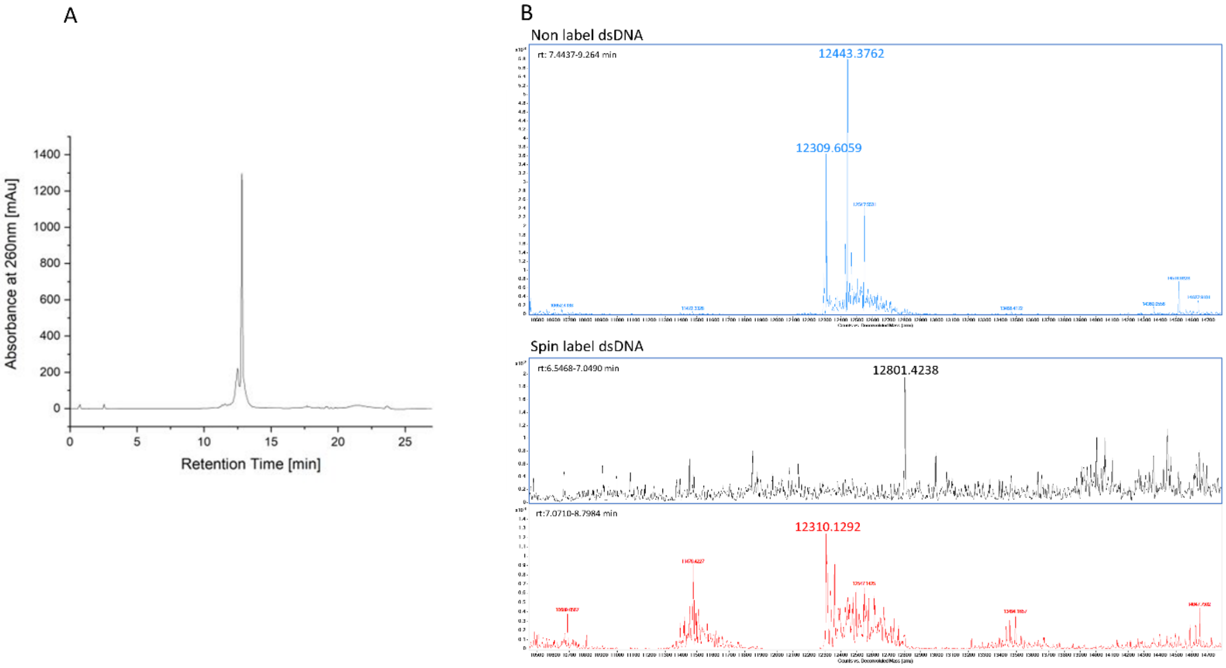


**Figure S2**: Characterization of representative spin-labeled DNA. **(A)** HPLC chromatogram measured at 260 nm of purified labeled *distal_Cop 10c*. **(B)** Deconvoluted mass spectra of non-labeled double-strand *distal_Cop 10c* (Mass of one strand is 12310 Da and the second is 12443 Da) (top) and spin-labeled *distal_Cop 10c* of each strand (labeled strand = 12801 Da, and non-labeled strand 12310 Da) (bottom). Similar HPLC purification profiles and mass spectrometry data were obtained for all DNA constructs, indicating successful labeling.

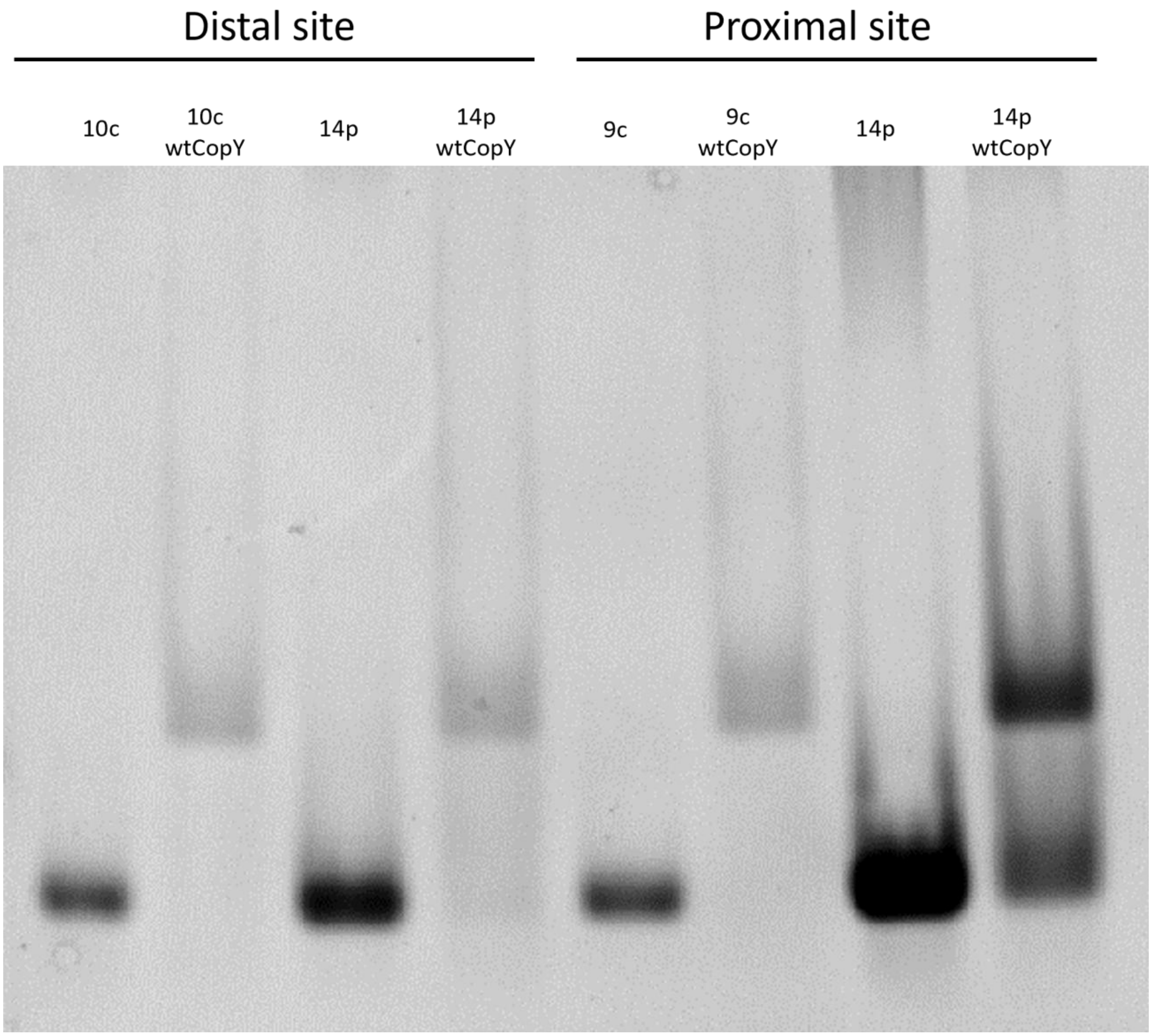

**Figure S3**: EMSA gel of WT-CopY binding to the various spin-labeled DNA.

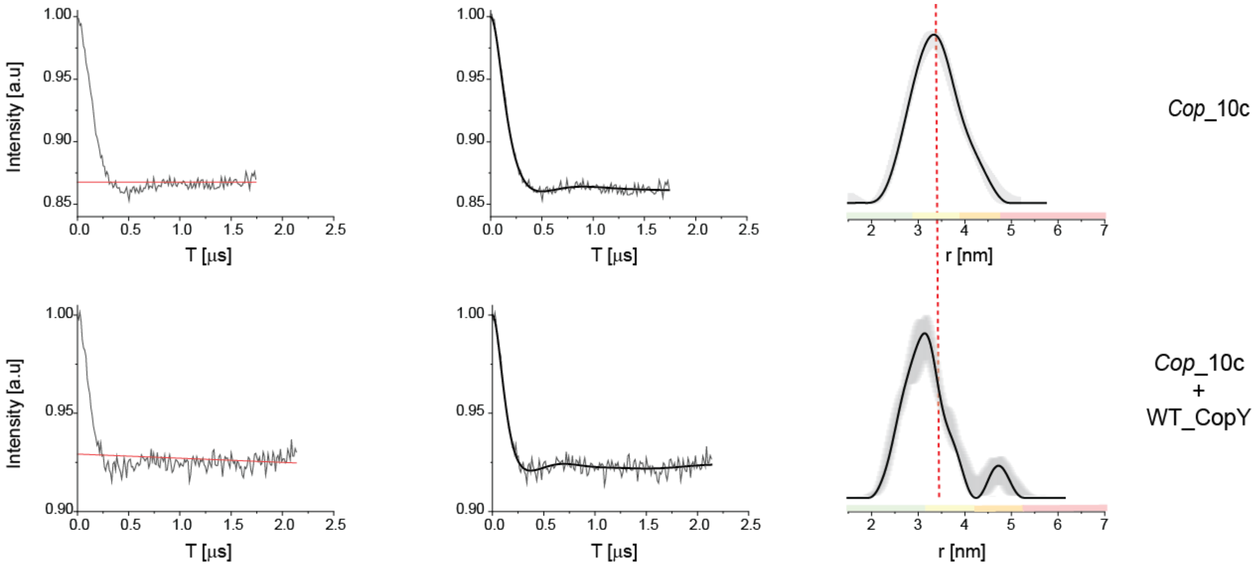


**Figure S5**: DEER on spin-labelled DNA distal_*Cop 10c*: Q-band DEER time domain signals before and after background subtraction and the corresponding distance distribution function. The data was analyzed using the DeerAnalysis program using Tikhonov regularization, where the regularization parameter was 20 (solid black lines). Distance distribution validation considered white noise, background start and dimensionality. The color bar indicates reliability ranges (green: shape reliable; yellow: mean and width reliable; orange: mean reliable; pink: no quantification possible).

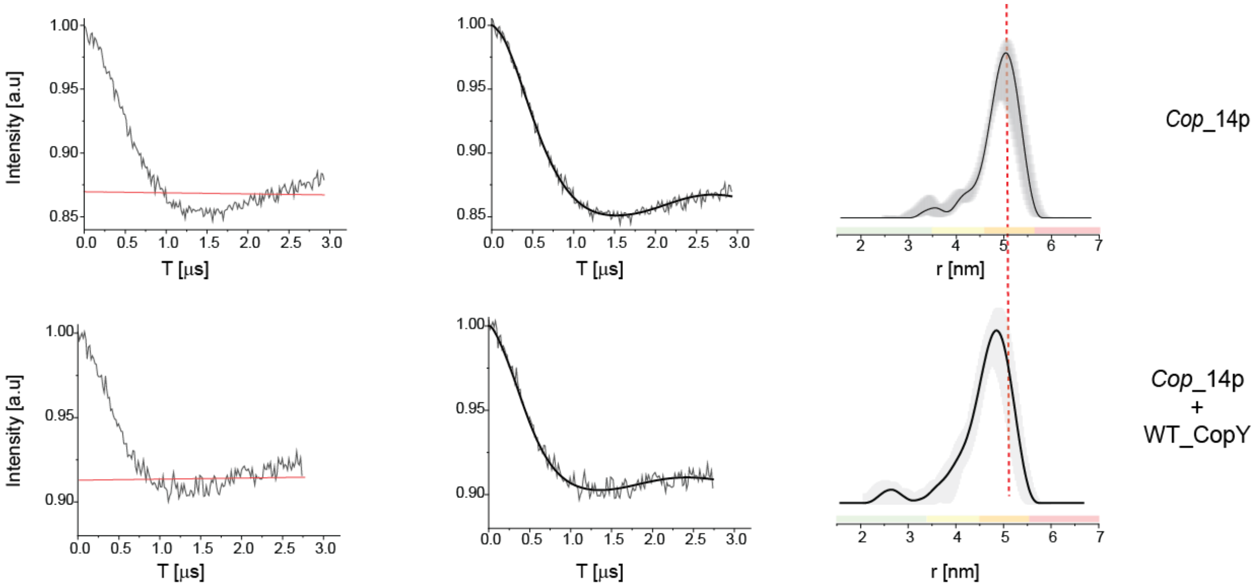


**Figure S5**: DEER on spin-labelled DNA distal_*Cop 14p*: Q-band DEER time domain signals before and after background subtraction and the corresponding distance distribution function. The data was analyzed using the DeerAnalysis program using Tikhonov regularization, where the regularization parameter was 20 (solid black lines). Distance distribution validation considered white noise, background start and dimensionality. The color bar indicates reliability ranges (green: shape reliable; yellow: mean and width reliable; orange: mean reliable; pink: no quantification possible).

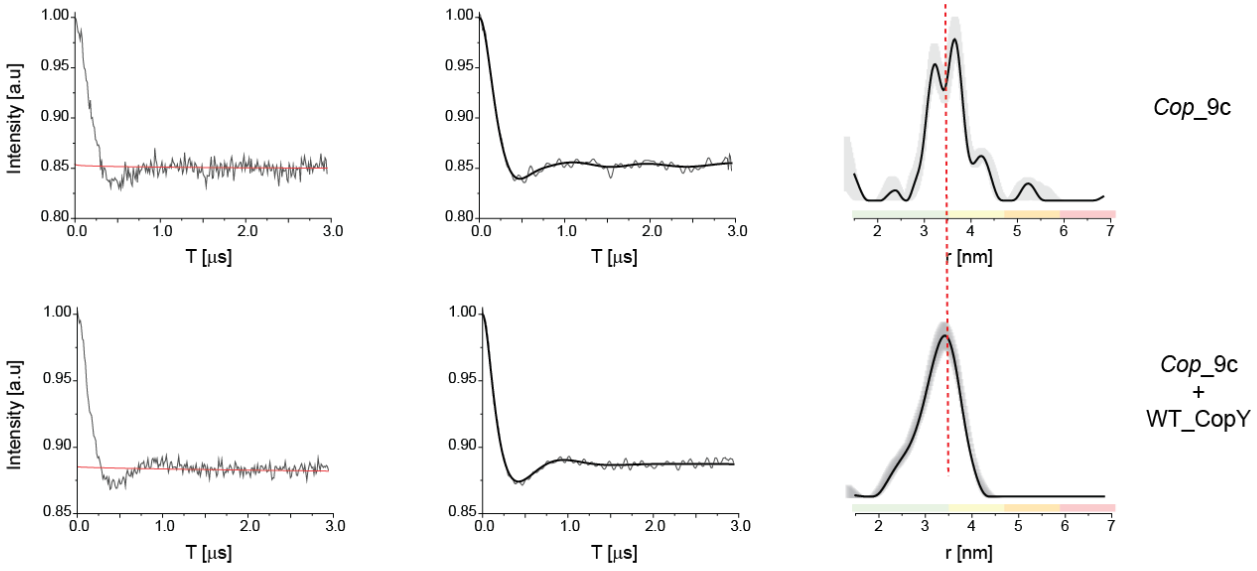


**Figure S6**: DEER on spin-labelled DNA proximal_*Cop 9c*: Q-band DEER time domain signals before and after background subtraction and the corresponding distance distribution function. The data was analyzed using the DeerAnalysis program using Tikhonov regularization, where the regularization parameter was 20 (solid black lines). Distance distribution validation considered white noise, background start and dimensionality. The color bar indicates reliability ranges (green: shape reliable; yellow: mean and width reliable; orange: mean reliable; pink: no quantification possible).

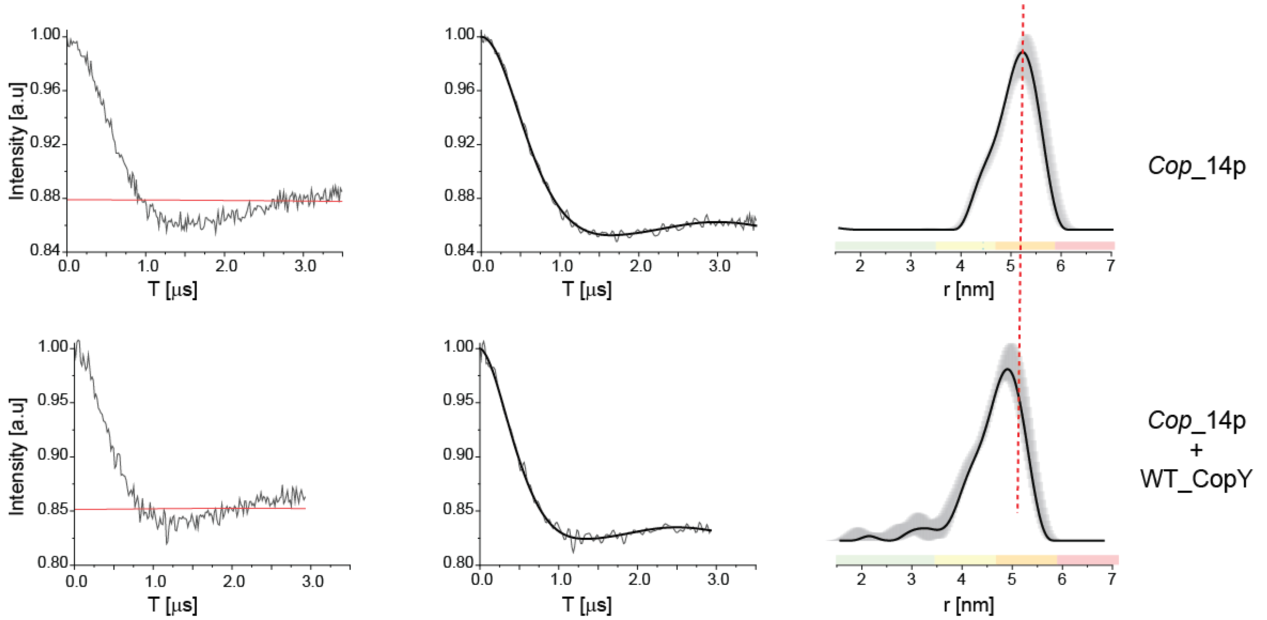


**Figure S7**: DEER on spin-labelled DNA proximal_*Cop 14p*: Q-band DEER time domain signals before and after background subtraction and the corresponding distance distribution function. The data was analyzed using the DeerAnalysis program using Tikhonov regularization, where the regularization parameter was 20 (solid black lines). Distance distribution validation considered white noise, background start and dimensionality. The color bar indicates reliability ranges (green: shape reliable; yellow: mean and width reliable; orange: mean reliable; pink: no quantification possible).

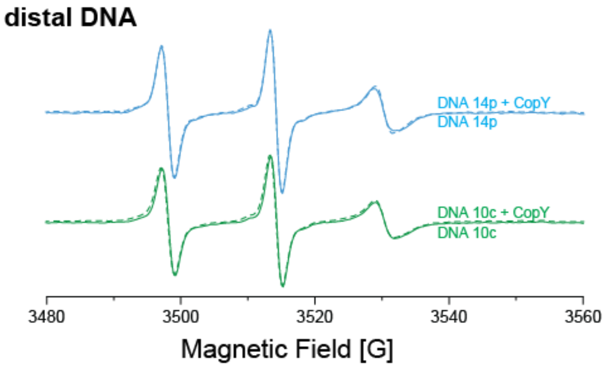


**Figure S8:** RT CW-EPR spectra for spin-labeled DNA in the absence (solid lines) and presence of CopY binding (dashed lines).

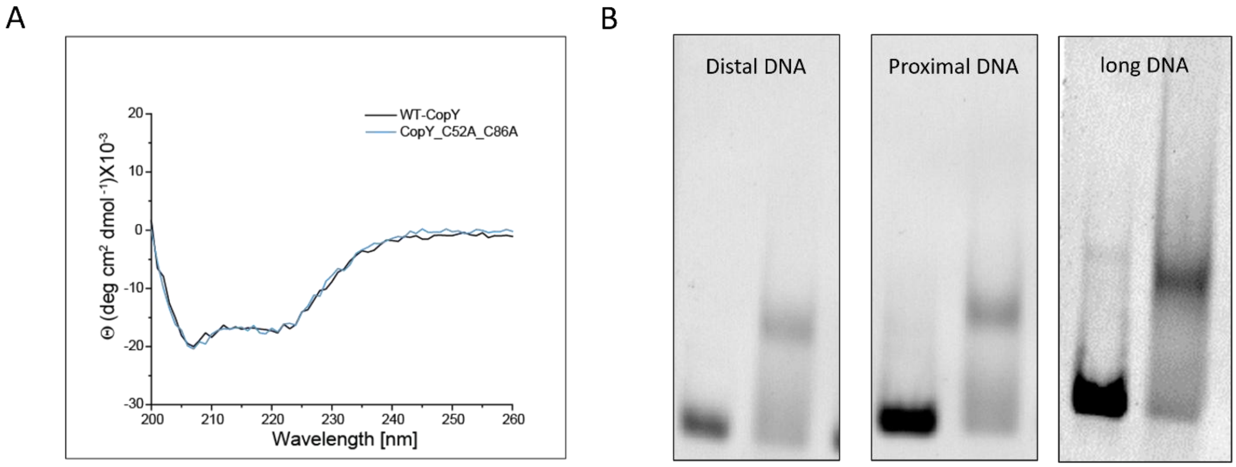


**Figure S9**: **A**. CD spectra of WT-CopY and CopY_C52A_C86A mutant. **B**. EMSA gel of CopY_C52A_C86A binding to the various DNAs.

**CD thermal analysis**

The thermal CD data were analyzed to compare the apparent thermal stability of CopY alone and in complex with long, distal, or proximal DNA. The buffer spectrum had already been subtracted during data acquisition. For each temperature, the CD signal was averaged between 220 and 224 nm to reduce single-wavelength noise. The common temperature range of 20-85 °C was used.

***Model-free observed T50***

Each trace was normalized using its measured values at 20 and 85 °C:

$$f_{obs}(T) = \frac{CD(T) - CD(20°\mathrm{C})}{CD(85°\mathrm{C}) - \mathrm{CD}(20°\mathrm{C})}$$

The observed $T_{50}$ was defined as the temperature at which $f_{obs}$ = 0.5 and was obtained by linear interpolation between the two surrounding experimental points. This analysis does not assume a thermodynamic model or require a post-transition plateau. It describes the midpoint of the change in the ellipticity between 20 and 85°C.

***Global van't Hoff fit***

A second analysis fitted the four non-normalized traces simultaneously. The van't Hoff relationship describes how the equilibrium between native (N) and unfolded (U) states changes with temperature. Under an apparent reversible two-state approximation, the predicted unfolded fraction is:

$$f_U(T) = \frac{1}{1 + exp\left[\frac{\Delta H_{vH}}{R}\left(\frac{1}{T_k} - \frac{1}{T_{m,k}}\right)\right]}$$

Here, $T_K$ is the temperature in Kelvin. R (gas constant) = 0.00831446 kJ $mol^{-1}$ $K^{-1}$. At T = $T_m$, the exponent is zero and $f_U$ = 0.5. The parameter $\Delta H_{vH}$ determines the steepness of the transition: a larger value produces a sharper transition. The unfolded fraction was converted into a predicted CD signal using a linear baseline and a fitted transition amplitude:

$$CD_{fit}(T) = y_0 + m(T - T_m) + A\, f_U(T)$$

T is the experimental temperature. CD(T) is the mean CD signal at 220-224 nm. $y_0$ is vertical position of the native-state baseline near Tm. m is linear temperature dependence of the baseline. A is the signal change associated with the transition. $T_m$ is temperature at which the model predicts $f_U$ = 0.5. $\Delta H_{vH}$ is van't Hoff enthalpy. The value 110 kJ $mol^{-1}$ was used only as the starting estimate supplied to Origin program. During least-squares optimization, Origin adjusted the shared parameter and obtained $\Delta H_{vH}$ = 107 ± 17 kJ $mol^{-1}$.
